# Droplet single-cell CRISPR screens identify regulators of T cell–mediated target-cell killing

**DOI:** 10.64898/2026.06.23.734054

**Authors:** Giulia Saronio, Gaia Antonini, Ankit Jain, Ian Vogel, Rachele Teneggi, Julia Neumann, Giada Zoppi, Yun Ding, Matteo Pecoraro, Mathias Langner, Peter Mirtschink, Stavros Stavrakis, Andrew de Mello, Roger Geiger

## Abstract

Cytotoxic CD8⁺ T cells kill target cells through brief cell–cell encounters, but pooled genetic screens cannot readily link perturbations in individual T cells to the fate of the target cells they engage. We developed droplet single-cell CRISPR screening to pair individual primary human CD8⁺ T cells with cancer cells, measure rapid target-cell death, and recover sgRNAs from phenotype-defined droplets. Applied across primary T cells from multiple donors, the platform recovered regulators of T cell receptor signaling, synapse formation, granule exocytosis and cytotoxic differentiation, and identified negative regulators of killing, including established inhibitory nodes such as *PTEN*, *RASA2* and *FOXO1*, together with *AFAP1L2* and components of the mTORC1 pathway. Validation across bispecific engager and TCR-engineered settings showed that selected hits modulate target-cell killing across recognition modalities and tumor models. Unexpectedly, perturbation of *RPTOR* or *RHEB* enhanced cytotoxic execution while reducing mTORC1 output, increasing AKT phosphorylation, and attenuating anabolic programs. Transient pharmacologic mTORC1 inhibition reproduced this rapid-killing state and improved antitumor activity after adoptive transfer. These results establish an interaction-resolved pooled genetic strategy for mapping cytotoxicity regulators and reveal that transient modulation of mTORC1 can shift T cells from anabolic growth toward rapid cytotoxic execution to enhance antitumor activity.

## Introduction

Cytotoxic CD8⁺ T cells eliminate infected and malignant cells by forming immunological synapses and delivering lytic granules through polarized exocytosis ^1–4^. This contact-dependent process is central to endogenous antitumor immunity and engineered T cell therapies, yet the molecular pathways that determine how efficiently individual T cells kill target cells remain incompletely defined. In particular, genetic regulators of target-cell killing are difficult to resolve using assays that measure population-level tumor control or long-term T cell expansion.

A major obstacle to dissecting cytotoxicity at scale is that the phenotype of interest is not cell-autonomous. In contrast to proliferation, survival, or transcriptional output within the perturbed cell itself, T cell cytotoxicity is defined by the fate of a physically interacting target cell. Conventional pooled CRISPR screening formats are therefore poorly suited to killing phenotypes, because the perturbation is encoded in the T cell whereas the functional readout is death or survival of its paired target. Maintaining this linkage generally requires physical compartmentalization of individual effector–target pairs, a requirement that has typically favored low-throughput, arrayed screening approaches involving individual sgRNA reagents and multiwell functional assays ^5,6^.

Droplet microfluidics offers a way to compartmentalize cellular interactions at high throughput, and droplet-based genetic screens have begun to enable pooled analysis of multicellular communication ^7,8^. However, cytotoxic T cell killing presents a distinct functional challenge: the relevant output is target-cell death after a short-lived, contact-dependent encounter with an individual effector cell. We therefore set out to adapt droplet-based screening to primary human CD8⁺ T cells, with the goal of measuring target-cell killing while preserving the linkage between each T cell’s sgRNA and the viability outcome of its paired target cell.

Here, we describe a droplet single-cell CRISPR screening platform that enables pooled functional genomics of primary human T cell cytotoxicity. The assay pairs individual genetically perturbed CD8⁺ T cells with cancer targets, quantifies target-cell death over short time windows, and recovers sgRNAs from droplets selected by target-cell fate. Using this approach, we recovered known regulators of T cell receptor signaling, synapse formation, granule exocytosis and cytotoxic differentiation, and identified previously unrecognized negative regulators of killing, including *AFAP1L2* and components of the mTORC1 pathway. Follow-up studies showed that perturbation of *RPTOR* or *RHEB* accelerated early per-cell killing while reducing anabolic programs and proliferative fitness, illustrating how cytotoxicity can be uncoupled from longer-term growth. Together, these results establish an interaction-resolved screening strategy for mapping regulators of T cell–mediated target-cell killing in primary human immune cells.

## Results

### A droplet assay measures rapid T cell cytotoxicity at single-cell resolution

We established a droplet-based microfluidic workflow to measure T cell-mediated cytotoxicity at single-cell resolution. Primary human CD8⁺ T cells and CD19⁺ JeKo-1 lymphoma target cells were co-encapsulated in picoliter water-in-oil droplets together with the CD19 × CD3 bispecific engager blinatumomab and the viability dye SYTOX Green (Fig. 1a). Droplets were generated at room temperature and incubated at 37 °C for defined periods to allow T cell-mediated target-cell killing.

**Figure 1.**
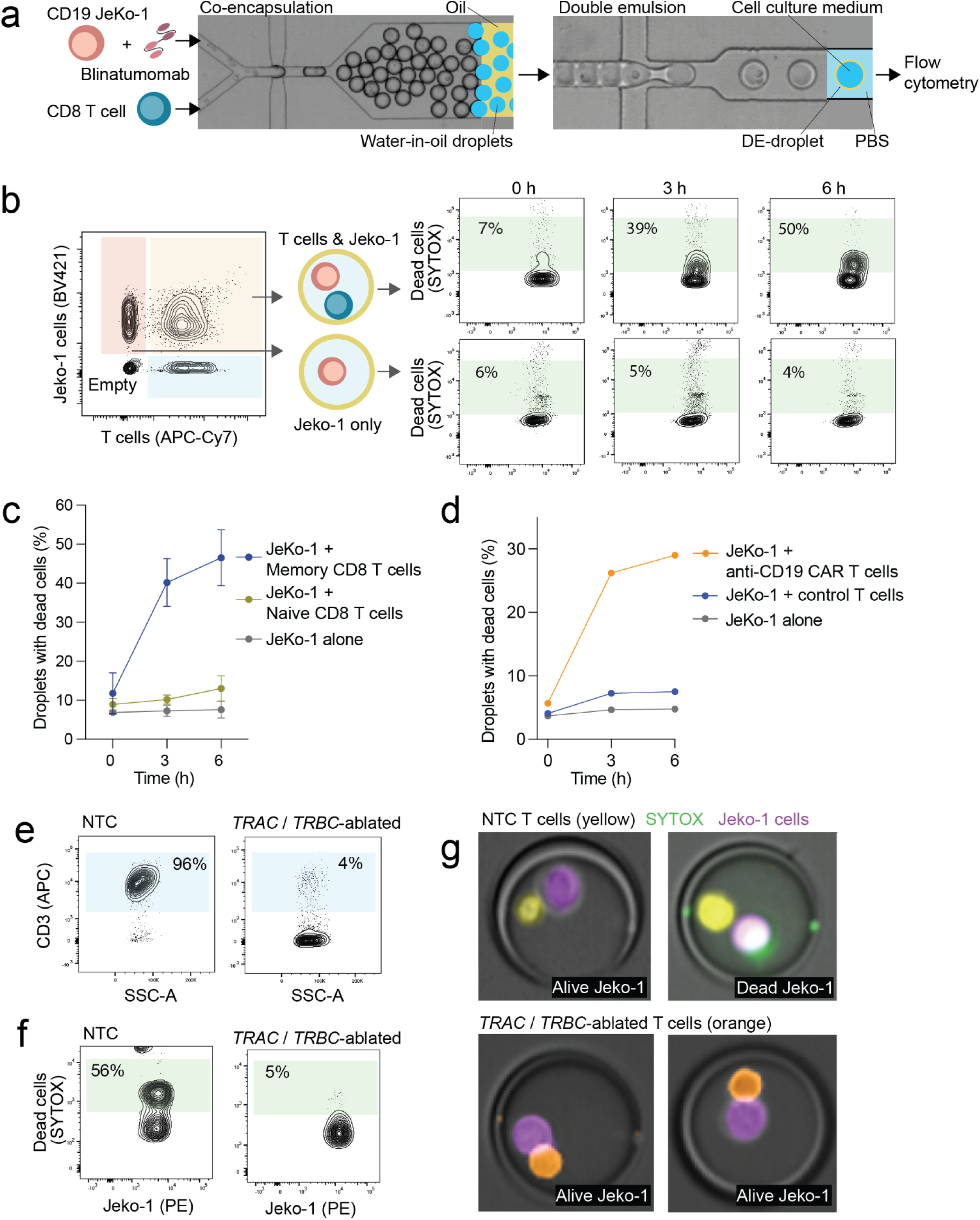
A droplet microfluidic platform quantifies rapid single-cell CTL-mediated killing. **(a)** Schematic of the assay workflow. Primary human CD8⁺ T cells and CD19⁺ JeKo-1 lymphoma cells were co-encapsulated with blinatumomab and SYTOX Green in water-in-oil droplets using a microfluidic device. Droplets were subsequently converted to double emulsions and analyzed by conventional flow cytometry. **(b)** Representative flow-cytometry analysis of double-emulsion droplets. Droplets containing both a T cell and a JeKo-1 cell were distinguished from droplets containing JeKo-1 cells alone, and target-cell death was quantified by SYTOX Green positivity at the indicated time points. Percentages indicate dead-target droplets within each gated population. **(c)** Time course of JeKo-1 killing by FACS-purified memory or naïve CD8⁺ T cells in droplets. JeKo-1 cells alone served as a viability control. Data are mean ± SEM; n = 2 donors. **(d)** Time course of JeKo-1 killing by anti-CD19 CAR-T cells compared with control T cells in droplets. JeKo-1 cells alone served as a viability control. **(e)** Flow-cytometric validation of CRISPR/Cas9-mediated disruption of TRAC and TRBC in memory CD8⁺ T cells. Surface CD3 expression is shown for non-targeting control (NTC) and TRAC/TRBC-ablated cells. **(f)** Representative droplet killing assay comparing NTC and TRAC/TRBC-ablated T cells co-encapsulated with JeKo-1 cells. Target-cell death was quantified by SYTOX Green positivity. Percentages indicate dead-target droplets. **(g)** Representative imaging flow-cytometry images of droplets containing T cells and JeKo-1 cells. NTC T cells formed productive lytic contacts associated with target-cell death, whereas TRAC/TRBC-ablated T cells formed non-lytic contacts and failed to induce death.

To enable quantitative phenotypic analysis by conventional flow cytometry, water-in-oil droplets were converted into double emulsions comprising an aqueous core surrounded by a thin oil shell ^9^. These double-emulsion droplets remained stable in PBS for at least 24 h and were compatible with standard cytometer fluidics. This format enabled side-by-side flow-cytometric gating of droplets containing both a T cell and a target cell versus target-only droplets, and allowed target-cell death quantification within each droplet population (Fig. 1b).

Using this workflow, we observed a time-dependent increase in target-cell death specifically within droplets containing both a CD8⁺ T cell and a JeKo-1 cell, with a half-time of approximately 60-90 min (Fig. 1b). In contrast, control droplets containing only T cells or only JeKo-1 cells showed minimal cell death for at least 6 h (Supplementary Fig. 1). These data show that the assay captures rapid, T cell-dependent target-cell killing over a dynamic range.

### Validation with defined T cell subsets and targeting modalities

To determine whether the assay could resolve expected differences in cytotoxic capacity between T cell subsets, we compared FACS-purified naïve and memory CD8⁺ T cells. Memory CD8⁺ T cells efficiently killed JeKo-1 cells, whereas naïve CD8⁺ T cells showed little or no cytotoxic activity under the same conditions (Fig. 1c), consistent with known functional differences between these subsets ^10^.

We next tested whether the platform could measure killing triggered by different modes of target-cell recognition. Anti-CD19 CAR-T cells killed JeKo-1 cells with kinetics similar to those observed for blinatumomab-activated memory CD8⁺ T cells, whereas control T cells lacking CD19-directed recognition showed no detectable cytotoxicity against JeKo-1 cells (Fig. 1d). Thus, the assay measures rapid target-cell killing mediated either by a soluble T cell engager or by a transgenic CAR.

To further validate that killing in droplets depended on CD3/TCR activation, we disrupted the TCRα and TCRβ constant loci, *TRAC* and *TRBC*, in memory CD8⁺ T cells using CRISPR/Cas9. Flow cytometry confirmed efficient loss of surface TCR/CD3 expression, with more than 90% of cells becoming TCR-negative (Fig. 1e). As expected, TCR-deficient T cells failed to kill JeKo-1 cells in droplets (Fig. 1f).

Imaging flow cytometry further confirmed that target-cell death required CD3/TCR triggering. Control T cells formed lytic contacts associated with target-cell death, whereas TCR-deficient T cells formed persistent but non-lytic contacts and failed to induce target-cell death (Fig. 1g). Together, these experiments establish that the droplet assay captures single-cell CTL killing in a specific and screening-compatible format.

### Pooled droplet CRISPR screening identifies regulators of rapid T cell killing

Having established a single-cell killing assay, we next adapted it for pooled CRISPR/Cas9 screening in primary human CD8⁺ T cells. Freshly isolated memory CD8⁺ T cells were activated and transduced with a custom sgRNA library comprising 4,462 sgRNAs targeting 791 genes involved in T cell signaling, differentiation, membrane trafficking and related cellular processes (Supplementary Table 1). After 48 h, cells were electroporated with recombinant Cas9 protein^11^ and cultured in puromycin to select sgRNA-expressing cells. Following 14 days of expansion, edited T cells were co-encapsulated with JeKo-1 target cells and blinatumomab in picoliter water-in-oil droplets containing SYTOX Green and incubated for 90 min at 37 °C to identify perturbations that impaired or enhanced rapid cytotoxicity (Fig. 2a).

**Figure 2.**
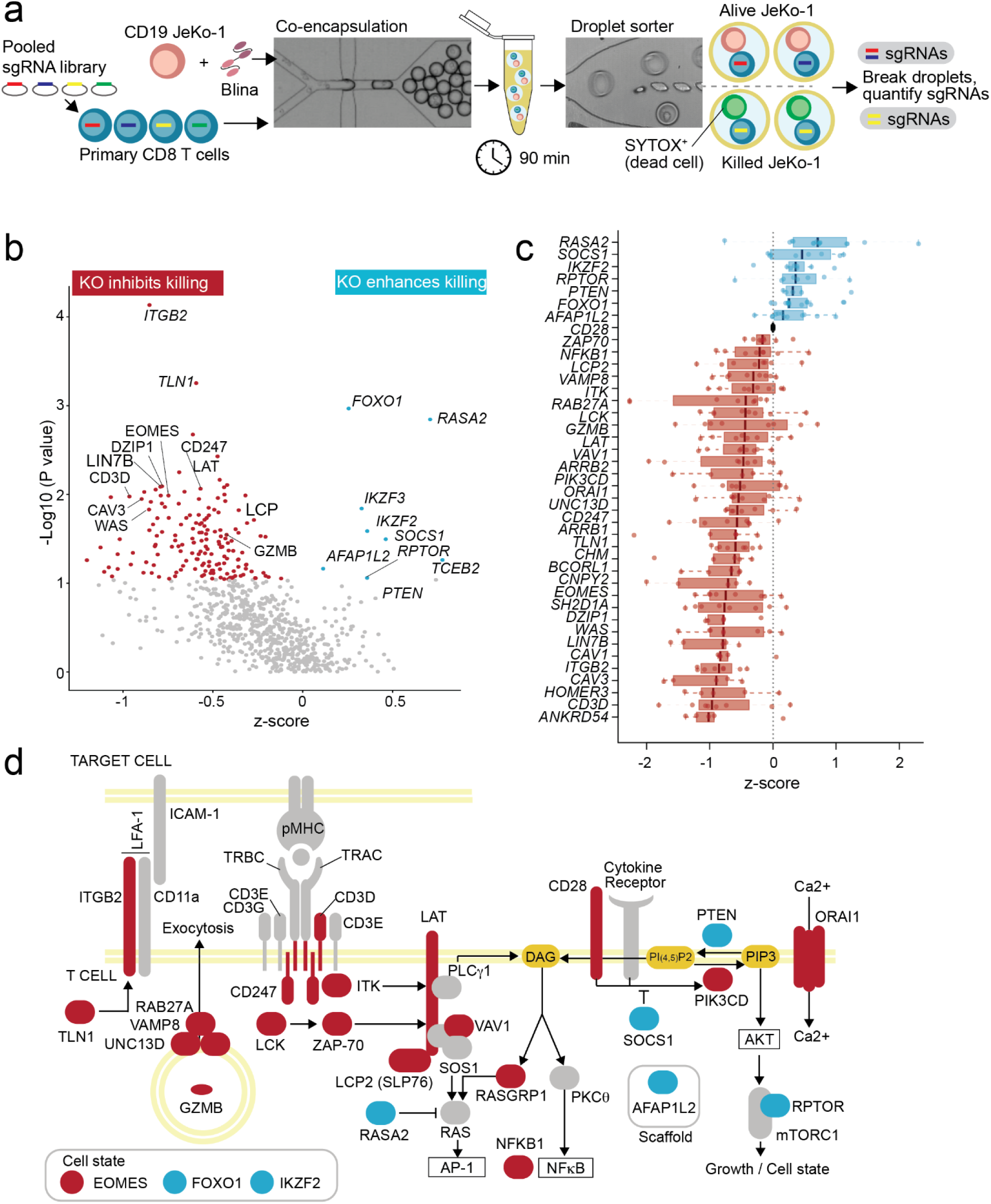
Pooled droplet CRISPR screening identifies regulators of rapid CTL-mediated killing. **(a)** Schematic of the pooled screening workflow. Primary human CD8⁺ T cells transduced with a pooled sgRNA library were co-encapsulated with JeKo-1 cells and blinatumomab. After 90 min, droplets containing killed targets (SYTOX⁺) or surviving targets were isolated by droplet sorting, broken, and subjected to sgRNA recovery and sequencing. **(b)** Gene-level screen summary. Each dot represents one targeted gene and shows the mean z-score from at least five independent screens, each performed with T cells from a different donor. Negative z-scores indicate genes whose disruption impaired rapid killing, whereas positive z-scores indicate genes whose disruption enhanced killing. Selected hits are labeled. **(c)** Distribution of gene-level effects for representative positive and negative regulators recovered in the screen. Points represent individual measurements with primary T cells from different donors across screens; boxes summarize the distribution for each gene. **(d)** Pathway map highlighting representative hits identified in the screen. Genes whose disruption impaired killing are shown as positive regulators of cytotoxicity, whereas genes whose disruption enhanced killing are shown as negative regulators of rapid killing.

To increase throughput, we bypassed double-emulsion generation at the sorting stage and used a custom high-speed water-in-oil droplet sorter operating at approximately 0.8 kHz ^12^. This workflow preserved sgRNA representation and recovery relative to flask-grown controls (R² = 0.93; Supplementary Fig. 2). To facilitate screening at scale, the sgRNA library was divided into three batches, each screened using primary CD8⁺ T cells from at least five healthy donors.

Each screen generated approximately 20 million droplets, of which roughly 15% contained both a T cell and a target cell. Among these droplets, approximately 25% exhibited target-cell killing after 90 min, corresponding to ∼750,000 informative events per screen. Droplets containing dead or live targets were sorted separately as target-dead and target-live fractions, broken and subjected to sgRNA recovery by PCR amplification and sequencing. Differential sgRNA representation between target-dead and target-live fractions was analyzed using MAGeCK ^13^, and gene-level statistics were aggregated across donors to identify perturbations that impaired or enhanced rapid killing (Fig. 2b,c). By linking each perturbed T cell to the viability outcome of its paired target cell over a short assay window, this workflow preferentially captures regulators of early killing kinetics rather than phenotypes dominated by proliferation or long-term outgrowth.

### The screen recovers known positive regulators of cytotoxicity

As expected for a loss-of-function screen of a coordinated effector process, perturbations more frequently impaired killing than enhanced it (Fig. 2b-d). Among these were multiple established positive regulators of CD8⁺ T cell cytotoxicity ^14,15^, validating the screening approach (Fig. 2b,c; Supplementary Table 2). These included components of the TCR complex and proximal signaling machinery, such as *CD3D*, *CD247*, *LCK*, *ZAP70*, *LAT*, *LCP2/SLP-76*, *VAV1*, and *ITK*, as well as the co-stimulatory receptor *CD28*, the PI3K pathway component *PIK3CD*, and the store-operated calcium channel *ORAI1*.

Beyond proximal T cell signaling, positive regulators spanned core modules of CTL-mediated target killing. These included lytic granule components and regulators of granule trafficking and exocytosis, such as *GZMB*, *UNC13D*, *RAB27A* and *VAMP8*; synapse and cytoskeletal regulators, including *ITGB2*, *TLN1* and *WAS*; and transcriptional regulators associated with CTL activation and cytotoxic differentiation, including *EOMES* and *NFKB1* ^16,17^. *SH2D1A* (SAP) also emerged as a positive regulator, consistent with its known role in SLAM-family receptor signaling and cytotoxic synapse formation against B-cell targets ^18^.

Beyond these expected hits, the screen identified several factors with less well-defined roles in CTL killing. These included *CAV1*, *ARRB1*, and *ARRB2*, which have been implicated in membrane organization and TCR redistribution at the synapse ^19,20^. Additional candidates, including *CAV3*, *HOMER3*, *CNPY2*, *LIN7B*, *DZIP1*, *CHM*, *BCORL1*, and *ANKRD54*, point to possible contributions from membrane organization, vesicle biology, cell polarity, and nuclear regulation. Collectively, these results indicate that the screen captures core features of CTL biology while identifying additional candidate regulators of rapid target-cell killing.

### The screen identifies negative regulators of rapid cytotoxicity

In addition to positive regulators, the screen identified genes whose disruption enhanced rapid target-cell killing within the 90-min assay window. These included established inhibitors of T cell activation, such as *RASA2*, *PTEN*, and *SOCS1* ^21–23^ (Fig. 2b,c; Supplementary Table 2). The screen also identified transcriptional regulators associated with quiescence or restricted effector differentiation, including *FOXO1* and *IKZF2* (Helios) ^24,25^. This suggests that programs maintaining quiescence or limiting effector differentiation can constrain the speed of target-cell killing.

Among the most notable hits were *AFAP1L2* and *RPTOR*, neither of which had been implicated in limiting rapid CTL cytotoxicity. *AFAP1L2* is a signaling adaptor enriched in dysfunctional tumor-infiltrating T cells ^26^, whereas *RPTOR* encodes Raptor, a defining component of mTORC1 ^27^. The emergence of *RPTOR* as a negative regulator was unexpected given the established role of mTORC1 in supporting T cell growth, metabolism, and effector differentiation ^28^, raising the possibility that the screen was capturing a distinct functional dimension of CTL biology that is not well resolved by proliferation-based or long-term assays.

### AFAP1L2 and RPTOR validate as negative regulators across assay formats

To validate *AFAP1L2* and *RPTOR* alongside established negative regulators, freshly isolated CD8⁺ T cells were edited by CRISPR/Cas9 and assayed in droplets together with JeKo-1 target cells and blinatumomab (Fig. 3a). Disruption of *AFAP1L2* or *RPTOR* consistently increased target-cell killing within the 90-min assay window, to an extent comparable to disruption of *RASA2*, *PTEN*, or *FOXO1* (Fig. 3b,c). Enhanced killing was also observed in orthogonal bulk co-culture assays measured after 8 h, confirming the phenotype across assay formats (Fig. 3d).

**Figure 3.**
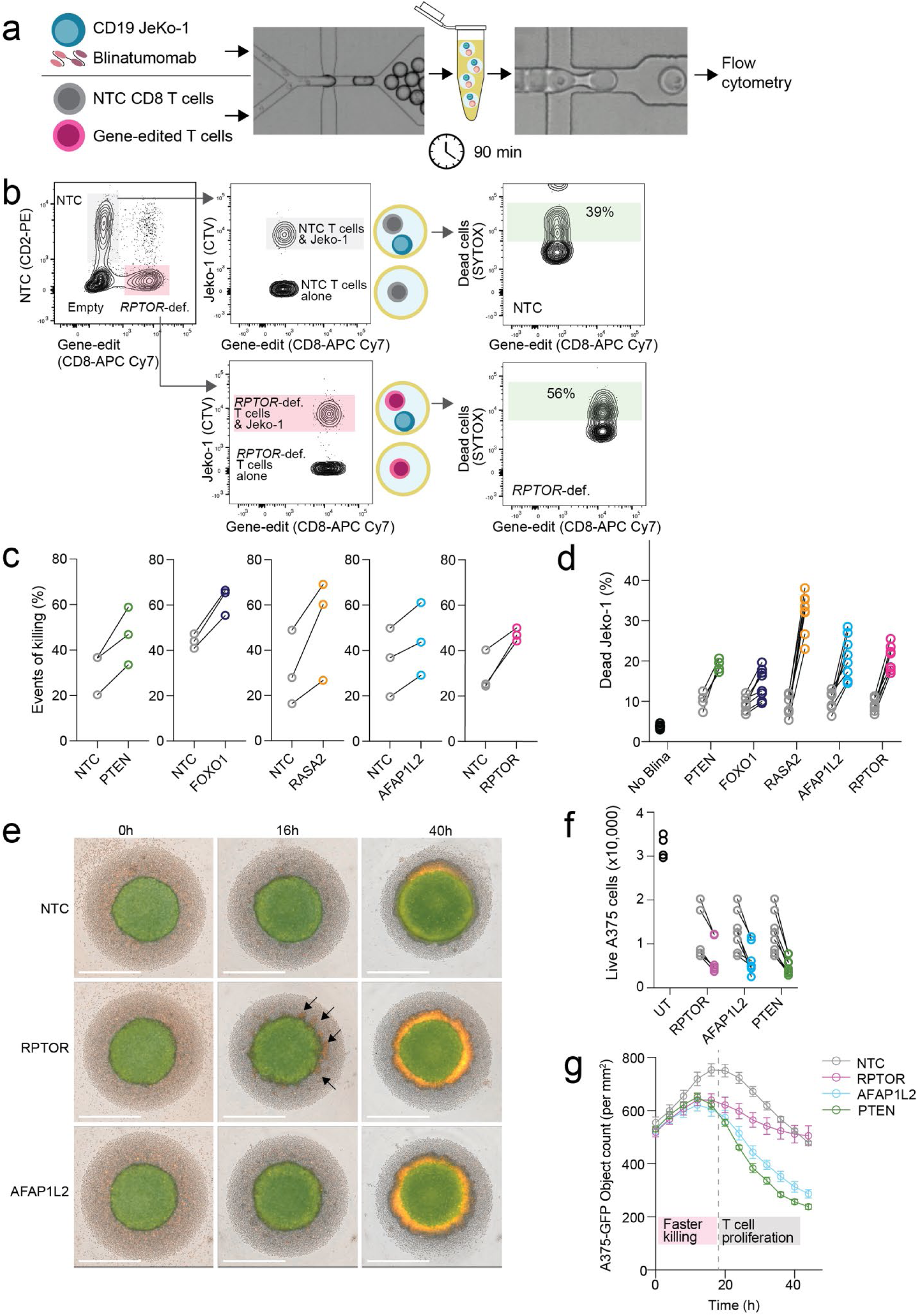
AFAP1L2 and RPTOR validate as negative regulators of rapid killing across assay formats and tumor models. **(a)** Schematic of the validation workflow. NTC and gene-edited primary human CD8⁺ T cells were differentially labeled, co-encapsulated with JeKo-1 cells and blinatumomab, incubated for 90 min, and analyzed by flow cytometry. **(b)** Representative gating strategy and example data from the competitive droplet assay. Fluorescence barcoding distinguished NTC from gene-edited T cells, gates identified droplets containing target cells, and target-cell death was quantified by SYTOX Green positivity. Representative plots for *RPTOR*-deficient cells are shown. **(c)** Quantification of droplet-based killing for the indicated gene disruptions. Loss of PTEN, FOXO1, RASA2, AFAP1L2, or RPTOR increased rapid JeKo-1 killing relative to matched NTC cells. Lines connect paired donor measurements. n=3 from three independent experiments with T cells from 3 different donors. **(d)** Orthogonal bulk co-culture validation assay. Percentage of dead JeKo-1 cells after co-culture with edited T cells is shown for the indicated perturbations; no-blinatumomab wells served as a negative control. **(e)** Representative images of tumor spheroid killing by 1G4 TCR-transduced T cells of the indicated genotypes at 0, 16, and 40 h. *RPTOR*- or *AFAP1L2*-edited T cells showed accelerated disruption and clearance of tumor spheroids compared with NTC cells. **(f)** Quantification of residual live A375 melanoma cells after co-culture with untransduced T cells (UT) or 1G4 TCR-transduced T cells of the indicated genotypes. **(g)** Kinetics of A375-GFP tumor-cell clearance by 1G4 TCR-transduced T cells. Early differences reflect faster killing by *RPTOR*-, *AFAP1L2*-, and *PTEN*-edited cells, whereas later time points capture the contribution of T cell proliferative fitness.

We next asked whether these findings extended beyond the CD19/blinatumomab system. Primary CD8⁺ T cells were transduced with the 1G4 TCR, which recognizes an NY-ESO-1-derived peptide presented by HLA-A*02:01 ^29^, and tested against solid tumor-derived target cells. In this setting, disruption of *RPTOR* or *AFAP1L2* accelerated killing of Huh7 spheroids and A375 melanoma cells relative to non-targeting control cells (Figs. 3e-g), indicating that both genes constrain short-term cytotoxic activity across antigen-recognition systems and cancer cell models.

At later time points, when cancer cell control became increasingly dependent on sustained T cell expansion, *RPTOR*-edited cells performed less well than control cells (Fig. 3g), consistent with the established role of mTORC1 in supporting T cell proliferation ^30^. These observations suggest that *RPTOR* disruption enhances rapid per-cell killing while compromising longer-term proliferative fitness, pointing to a functional separation between early cytotoxic execution and growth-related programs.

### RPTOR loss rewires mTORC1–AKT/MAPK signaling and enhances rapid cytotoxicity while attenuating anabolic programs

To determine whether the enhanced killing phenotype reflected impaired mTORC1 signaling rather than mTORC1-independent functions of Raptor, we ablated *RHEB*, a key upstream activator of mTORC1 ^27,31^. *RHEB*-deficient T cells also showed enhanced JeKo-1 killing, suggesting that reduced mTORC1 signaling underlies the rapid-killing phenotype observed after *RPTOR* perturbation (Fig. 4a).

**Figure 4.**
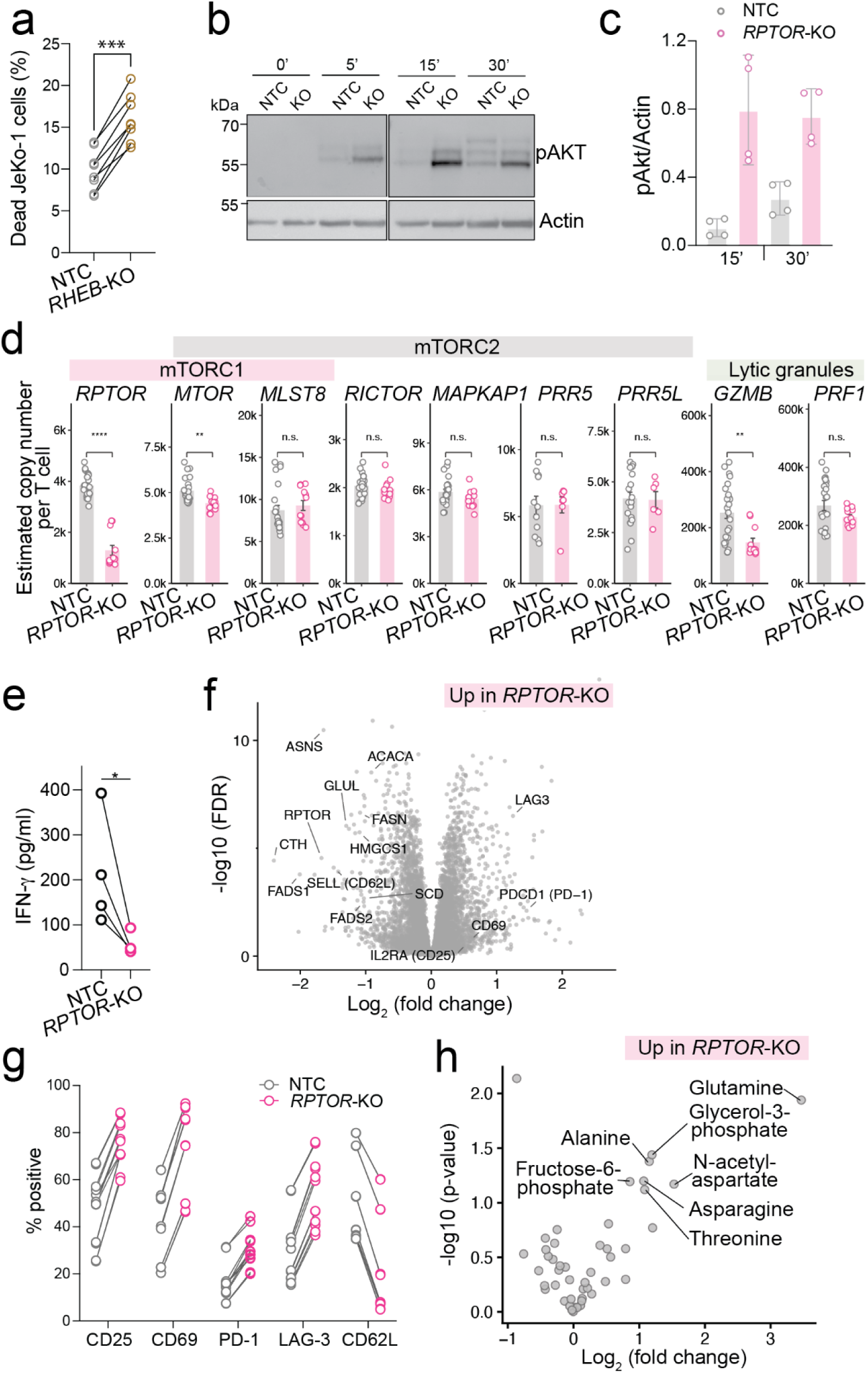
Reduced mTORC1 signaling enhances rapid cytotoxicity while restraining anabolic programs. **(a)** Validation of RHEB as a negative regulator of rapid cytotoxicity. *RHEB*-deficient T cells were assayed for JeKo-1 killing in bulk for 8h and showed enhanced target-cell killing relative to control cells. n=8, T cells from 4 donors measured in duplicates. Significance was determined by two-sided unpaired Student’s t-test. **(b)** Representative immunoblot of phospho-AKT (Ser473) in NTC and *RPTOR*-KO CD8⁺ T cells following anti-CD3/CD28 stimulation for the indicated times. Actin served as a loading control. **(c)** Densitometric quantification of phospho-AKT normalized to actin at 15 and 30 min after stimulation. Data are from two independent donors, each analyzed in duplicate. Bars indicate mean ± SEM. **(d)** Estimated copy numbers of mTOR complex subunits and cytotoxic effector proteins in resting NTC and *RPTOR*-KO T cells. Copy numbers per cell were derived from quantitative proteomics. RPTOR abundance was selectively reduced in *RPTOR*-KO cells, whereas mTORC2 subunits were unchanged; granule proteins are also shown. Mean ± SEM. (n=12 for *RPTOR*-deficient T cells from 4 donors, measured in triplicates), (n=24 for NTC T cells from 8 donors, measured in triplicates). Statistical comparisons were performed using two-sided Welch’s t-tests with Benjamini-Hochberg correction. **(e)** IFN-γ secretion by resting NTC and *RPTOR*-KO T cells measured by ELISA. Each point represents one independent donor; statistics by two-sided Welch-corrected t-test. **(f)** Differential protein abundance in resting *RPTOR*-KO versus NTC T cells by quantitative proteomics. Selected proteins with altered abundance are labeled, highlighting reduced anabolic and biosynthetic programs together with altered expression of cell-state markers. **(g)** Flow-cytometric analysis of activation and differentiation markers in resting NTC and *RPTOR*-KO T cells. Frequencies of CD25⁺, CD69⁺, PD-1⁺, LAG-3⁺, and CD62L⁺ cells are shown for each donor. **(h)** Differential metabolite abundance in resting *RPTOR*-KO versus NTC T cells by targeted metabolomics. Selected metabolites increased in *RPTOR*-KO cells are labeled.

To further define the signaling state induced by *RPTOR* disruption, we measured phosphorylation of canonical mTORC1 targets in edited primary CD8⁺ T cells. *RPTOR*-deficient cells showed reduced phosphorylation of p70 S6 kinase at Thr389, confirming impaired mTORC1 output (Supplementary Fig. 3). In the same cells, AKT phosphorylation was increased (Figs. 4b,c). Thus, *RPTOR* loss did not induce a globally heightened activation state, but instead was associated with increased AKT Ser473 phosphorylation. Together with the enhanced killing observed after *RHEB* disruption, these data support the conclusion that reduced mTORC1 signaling is associated with a distinct cytotoxic state characterized by accelerated target-cell killing, increased AKT phosphorylation and attenuated anabolic/proliferative programs.

We next performed quantitative proteomics on resting *RPTOR*-deficient and non-targeting control (NTC) T cells 12 days after activation, using cells from four independent donors (Fig. 4d). Copy-number estimates confirmed an approximately 65% reduction in Raptor abundance, from ∼3,800 to ∼1,300 copies per cell, whereas Rictor and other mTORC2 components were unchanged, consistent with selective disruption of mTORC1.

Perforin-1 and granzyme B abundance, as well as interferon-γ secretion, were reduced (Figs. 4d,e), arguing against increased lytic granule content as the basis for enhanced killing. Instead, the proteomic data pointed to a broader change in T cell state. SELL (CD62L) was among the most strongly depleted proteins, whereas LAG3 and PDCD1 (PD-1) were increased (Fig. 4f), suggesting that rested *RPTOR*-edited T cells adopt a more differentiated, effector-like phenotype. IL2RA (CD25) and CD69 were likewise elevated. These changes were confirmed by flow cytometry (Fig. 4g).

In parallel, *RPTOR* loss was associated with broad attenuation of anabolic programs. Enzymes involved in amino acid biosynthesis, including ASNS, GLUL, and CTH, were reduced, as were multiple enzymes involved in lipid synthesis, including FASN, ACACA, HMGCS1, FADS1, FADS2, and SCD (Fig. 4f). These data indicate an overall reduction in biosynthetic activity. Consistent with this interpretation, targeted metabolomics revealed accumulation of several amino acids and glycolytic intermediates in rested *RPTOR*-KO cells, suggesting reduced utilization of these metabolites for anabolic processes (Fig. 4h).

Together, these data indicate that reduced mTORC1 signaling enhances rapid target-cell killing without increasing lytic effector protein abundance and instead acts in the context of a broader reconfiguration of T cell state. Thus, rapid cytotoxic execution can be separated from anabolic and proliferative programs that are typically associated with T cell fitness.

### Transient mTORC1 inhibition enhances rapid killing and antitumor activity

The reduced proliferative fitness of *RPTOR*-edited T cells raised the question of whether enhanced rapid killing could be induced transiently, without permanent disruption of mTORC1 signaling. To test this, we treated activated CD8⁺ T cells with rapamycin, an inhibitor of mTORC1, for 8 days during *ex vivo* expansion. Similar to *RPTOR*-deficient T cells, rapamycin-treated CD8⁺ T cells killed JeKo-1 cells more rapidly in vitro (Fig. 5a,b).

**Figure 5.**
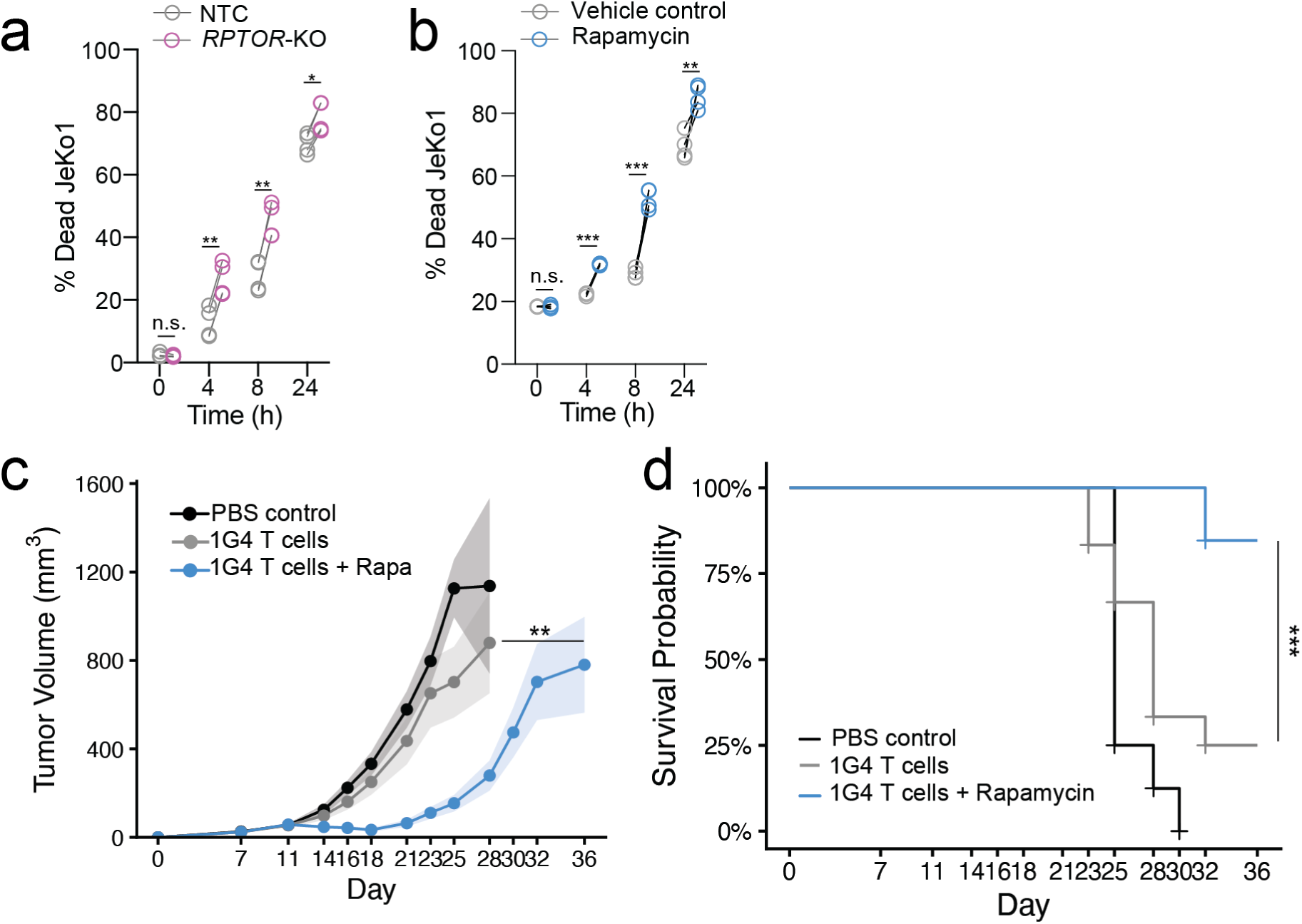
Transient mTORC1 inhibition improves antitumor activity in vivo. **(a)** Time course of JeKo-1 killing by NTC and RPTOR-edited T cells. *RPTOR* loss accelerated target-cell killing at early time points. Data are mean ± SEM (n = 4 per group from 2 different donors). Significance at each time point was determined by two-sided unpaired Student’s t-test. **(b)** Time course of JeKo-1 killing by vehicle- and rapamycin-treated T cells. Transient mTORC1 inhibition accelerated target-cell killing with kinetics similar to those observed after *RPTOR* disruption. Data are mean ± SEM (n = 4 per group from 2 donors). Significance at each time point was determined by two-sided unpaired Student’s t-test. **(c)** Tumor growth in A375 tumor-bearing NSG mice treated with PBS, 1G4 TCR-transduced T cells, or 1G4 TCR-transduced T cells manufactured in the presence of rapamycin. Data are mean tumor volume ± SEM; n = 10 (PBS), 12 (1G4), and 13 (1G4 + rapamycin). Statistical analysis was performed by two-way ANOVA. One representative of two independent experiments with similar results is shown. **(d)** Kaplan-Meier survival curves for mice in panel c. Tick marks indicate censored animals. Median survival was 25 days for PBS-treated mice, 28 days for 1G4-treated mice, and not reached for the 1G4 + rapamycin group. Statistical analysis was performed by log-rank test.

We next asked whether transient mTORC1 inhibition during ex vivo T cell expansion altered antitumor activity after adoptive transfer. 1G4 TCR-transduced T cells were treated with low-dose rapamycin during expansion and then transferred into A375 tumor-bearing NSG mice. Rapamycin-treated T cells improved tumor control and prolonged survival compared with vehicle-treated controls (Fig. 5c,d).

In contrast, T cells carrying permanent *RPTOR* or *RHEB* disruption did not improve tumor control *in vivo*, coinciding with their reduced proliferation rate (Supplementary Fig. 4). Thus, transient pharmacologic mTORC1 inhibition enhanced the rapid-killing phenotype without the apparent *in vivo* fitness cost associated with permanent genetic disruption of the pathway.

## Discussion

We developed a droplet single-cell CRISPR screening platform to genetically dissect T cell–mediated target-cell killing. By preserving the linkage between a perturbed T cell and the viability outcome of its paired target cell, the assay converts a transient, non-cell-autonomous immune effector function into a pooled genetic screening phenotype. This builds on prior droplet-based screening strategies for cell–cell communication ^7^, but applies compartmentalized screening to a native cytotoxic effector function in primary human T cells. By resolving individual effector–target-cell interactions over short time windows, the assay enables genetic analysis of cytotoxicity with kinetic resolution and complements CRISPR approaches that primarily capture proliferation, persistence or other longer-term cell-intrinsic states.

Applying this platform to primary human CD8⁺ T cells recovered canonical cytotoxicity pathways and identified regulators of target-cell killing, including *AFAP1L2* and *RPTOR*. Several hits mapped to expected modules of T cell receptor signaling, synapse formation and granule exocytosis, whereas others pointed to less well-characterized pathways that constrain cytotoxicity. The identification of *RPTOR* as a negative regulator was unexpected given the established role of mTORC1 in supporting T cell growth, metabolism and effector differentiation ^27,28^. However, prior work has shown that mTOR inhibition does not simply suppress CD8⁺ T cell function: rapamycin can promote memory differentiation and improve recall capacity, and mTOR signaling helps set the balance between effector and memory fate ^32^. Our data add a kinetic dimension to this framework by showing that reduced mTORC1 signaling can accelerate early per-cell target killing even as anabolic metabolism and proliferative fitness decline.

These findings suggest that cytotoxic execution and growth-associated T cell fitness can be uncoupled. Reduced mTORC1 signaling favored early target-cell killing but compromised sustained expansion when the pathway was permanently disrupted. Consistent with this, durable disruption of *RPTOR* or *RHEB* did not improve tumor control *in vivo*, whereas transient rapamycin exposure during *ex vivo* expansion enhanced antitumor activity after adoptive transfer. This distinction is important because previous studies have linked pharmacologic mTOR modulation to favorable CD8⁺ T cell states, including memory-like properties relevant to adoptive therapies ^33,34^. Our results connect this tunable pathway to a screen-derived cytotoxicity phenotype and distinguish transient pharmacologic modulation from permanent genetic loss of mTORC1 signaling.

The proximal mechanisms by which reduced mTORC1 signaling accelerates target-cell killing remain to be fully defined. Our phospho-signaling analysis suggests that *RPTOR* disruption shifts T cells into a state with reduced mTORC1 output and increased AKT phosphorylation. Increased AKT phosphorylation following mTORC1 inhibition is consistent with release of established mTORC1/S6K-dependent negative-feedback circuits that restrain upstream receptor–PI3K–AKT signaling in other cellular contexts^35^. However, the functional consequence of this feedback for CTL-mediated target-cell killing was not predictable from previous work. Reduced mTORC1 signaling appears to uncouple cytotoxic execution from anabolic growth programs, favoring rapid target-cell killing at the expense of sustained proliferative fitness.

More broadly, this study shows that interaction-resolved genetic screens can uncover regulators of immune-cell function that are not readily captured by conventional cell-autonomous assays. Although the current implementation focuses on T cell cytotoxicity, the same principle could be extended to other immunological interactions in which the key phenotype is encoded in the response of a cellular partner. Such approaches may help define genetic programs that control not only what immune cells become, but how they act on other cells.

## Acknowledgments

We thank members of the Geiger and deMello laboratories for helpful discussions and feedback on the manuscript. This work was supported by the Swiss Science Foundation 310030_197737 (RG), the European Research Council 803150 (RG), Swiss Cancer Research KFS-5537-02-2022 (RG), the Helmut Horten Foundation (AdM and RG) and the Institute for Research in Biomedicine.

## Author contributions

Conceptualization: GS, RG

Methodology: GS, GA, IV, AJ, RT, JN, GZ, YD, ML, MP

Investigation: GS, GA, IV, AJ, RT, JN, GZ, YD, ML, MP

Formal analysis: GS, GA, RG

Resources: AdM, RG

Supervision: PM, SS, AdM, RG

Funding acquisition: AdM, RG

Writing – original draft: GS, RG

Writing – review & editing: all authors

## Data and materials availability

The mass spectrometry proteomics data have been deposited to the ProteomeXchange Consortium via the PRIDE ^36^ partner repository with the dataset identifier PXD077938.

## Materials and Methods

### Cell culture, primary cells, and cell lines

Cells were cultured at 37 °C in a humidified incubator with 5% CO₂. Unless otherwise indicated, all cell culture reagents were purchased from Gibco (Thermo Fisher Scientific, USA). Complete RPMI medium (cRPMI) consisted of RPMI-1640 supplemented with 10% heat-inactivated fetal bovine serum (HI-FBS), 1× GlutaMAX, 1× penicillin/streptomycin, 1× kanamycin, 1× HEPES, 1× non-essential amino acids, 1× sodium pyruvate, and 50 μM β-mercaptoethanol. Complete DMEM (cDMEM) consisted of DMEM supplemented with 10% HI-FBS, 1× GlutaMAX, 1× penicillin/streptomycin, 1× kanamycin, 1× HEPES, 1× non-essential amino acids, and 1× sodium pyruvate. Complete Opti-MEM (cOpti-MEM) consisted of Opti-MEM supplemented with 5% HI-FBS, 1× non-essential amino acids, 1× sodium pyruvate, and 50 μM β-mercaptoethanol.

Peripheral blood from healthy donors was obtained from the Swiss Blood Donation Center of Lugano in accordance with approval from the Swiss Federal Office of Public Health (authorization 2018-02166/CE 3428). Peripheral blood mononuclear cells (PBMCs) were isolated by Ficoll-Paque density-gradient centrifugation (Sigma-Aldrich, Buchs, Switzerland). Primary human CD8⁺ and CD4⁺ T cells were enriched from PBMCs using magnetic microbeads (Miltenyi Biotec, Adliswil, Switzerland).

For experiments requiring purified subsets, CD8⁺ T cells were sorted on a SORP FACSymphony S6 cell sorter (BD Biosciences) based on CD45RA and CCR7 expression. Naïve CD8⁺ T cells were defined as CD45RA⁺CCR7⁺, central memory T cells as CD45RA⁻CCR7⁺, and effector memory T cells as CD45RA⁻CCR7⁻. Where indicated, memory CD8⁺ T cells were cultured in cRPMI supplemented with recombinant human IL-7 (10 ng/ml; Sino Biological) and IL-15 (10 ng/ml; Sino Biological).

JeKo-1 cells (ATCC CRL-3006) were maintained in cRPMI. Lenti-X 293T cells (Takara Bio, cat. no. 632180), A375 cells (ATCC CRL-1619) and Huh7 (JCRB0403) were cultured in cDMEM

### sgRNA library generation and molecular cloning

The sgRNA library was designed to target protein-coding genes involved in TCR signaling, cytoskeletal remodeling, granule biology, ciliogenesis, and cellular metabolism. Candidate guides were selected using the CRISPOR web tool and prioritized from the Brunello and Toronto KnockOut libraries. The final library comprised 4,462 sgRNAs targeting the selected genes, with 4–6 sgRNAs per gene, together with 100 non-targeting control sgRNAs. Oligonucleotide pools were synthesized by Twist Bioscience (South San Francisco, CA, USA).

The oligonucleotide pool was PCR-amplified using Phusion High-Fidelity PCR Master Mix (NEB, cat. no. M0531). BsmBI-v2 restriction sites and appropriate overhangs were introduced to enable cloning into the lenti-sgRNA puro backbone (Addgene plasmid 104990) by Golden Gate assembly (NEB, cat. no. E1601). The assembled product was electroporated into Lucigen Endura electrocompetent cells (Lucigen, cat. no. 60242-2). Following 1 h recovery in recovery medium (LGC Biosearch Technologies, cat. no. 80026-1), serial dilutions were plated to determine transformation efficiency and confirm adequate library coverage. The remaining culture was expanded overnight at 30 °C in 150 ml LB medium, and plasmid DNA was purified using a NucleoBond Xtra Midi kit (Macherey-Nagel, cat. no. 740410.50).

### Lentivirus production

Lentiviral particles were generated in Lenti-X 293T cells using the second-generation packaging plasmids psPAX2 (15 μg; Addgene #12260) and pMD2.G (5 μg; Addgene #12259) together with 20 μg transfer vector. Plasmids were diluted in Opti-MEM and incubated with PEI MAX (1 mg/ml; linear polyethylenimine hydrochloride, MW 40,000) for 10 min at room temperature before addition to Lenti-X 293T cells at approximately 80% confluency. Cells were incubated at 37 °C for at least 8 h, after which the medium was replaced with cOpti-MEM supplemented with 1× ViralBoost Reagent (ALSTEMBio, cat. no. VB100). Viral supernatants were collected the following day and centrifuged at 2,000 rpm for 5 min at 4 °C to remove cell debris. For concentration, cleared supernatants were mixed 3:1 with Lenti-X concentrator solution and incubated at 4 °C for 4 h, followed by centrifugation at 1,500 × g for 45 min at 4 °C. Pellets were resuspended in PBS at 1/100 of the original volume.

### T-cell activation, transduction, and gene editing Pooled screens

Primary human memory CD8⁺ T cells were activated in 96-well Nunc MaxiSorb plates (Thermo Fisher Scientific) coated with anti-CD3 (5 μg/ml; clone TR66, produced in-house) and anti-CD28 (1 μg/ml; clone CD28.2, BD Biosciences) at 2 × 10^6 cells per well in cRPMI. Twenty-four hours after stimulation, concentrated lenti-sgRNA puro virus was added at a multiplicity of infection (MOI) of 0.3, and cells were spin-infected at 800 × g for 45 min at 32 °C.

Twenty-four hours later, cells were collected, pelleted, and resuspended in 17 μl P3 buffer (Lonza, cat. no. V4XP-3032) per 10^6 cells. Cas9 ribonucleoprotein (RNP) complexes were added in a final volume of 3 μl per reaction. The 20 μl cell suspension was transferred to a 96-well Nucleocuvette plate (Lonza, cat. no. VVPA-1002) and electroporated on a 4D-Nucleofector (Lonza) using pulse code EH115. Immediately after electroporation, 80 μl antibiotic-free cRPMI was added to each well, and cells were incubated for 20 min at 37 °C before transfer to 24-well plates at 10^6 cells/ml. Two days later, puromycin (1.5 μg/ml; Sigma-Aldrich, cat. no. P8833) was added for 1 week to select sgRNA-expressing cells. Cells were expanded every 2 days with fresh medium while maintaining library representation.

### Arrayed knockout by RNP electroporation

Custom crRNAs targeting genes of interest, negative-control crRNA (IDT, cat. no. 1072544), and tracrRNA (IDT, cat. no. 1072532) were synthesized by IDT and resuspended in nuclease-free duplex buffer at 200 μM. To assemble guide duplexes, crRNA and tracrRNA were mixed 1:1 (v/v), heated to 95 °C for 5 min, and slowly cooled to room temperature. Recombinant Cas9 protein (Invitrogen, 5 mg/ml stock) was added at a 1:1 volume ratio relative to guide duplex and incubated for 20 min at room temperature. Poly(glutamic acid) (PGA; 100 mM) was added at 1.2 μl per 1.5 μl guide duplex to stabilize RNP formation.

Freshly isolated primary human T cells were activated for 48 h on plate-bound anti-CD3 and anti-CD28 as described above, collected, and resuspended in P3 buffer at 10^6 cells per 17 μl. Cells were mixed with 3 μl RNP complex, transferred to a 96-well electroporation plate, and electroporated using pulse code EH115. Immediately after electroporation, 80 μl pre-warmed antibiotic-free cRPMI was added to each well, and cells were incubated for 20 min at 37 °C. Cells were then transferred to 24-well plates at 10^6 cells/ml and allowed to recover for 2 days before expansion for an additional 7 days in cRPMI.

### Assessment of knockout efficiency

Genomic DNA was extracted using QuickExtract DNA Extraction Solution (Lucigen) according to the manufacturer’s instructions. Target loci were PCR-amplified from edited cells and from cells electroporated with a non-targeting control guide. PCR products were purified using a PCR clean-up kit (Macherey-Nagel) and submitted for Sanger sequencing (Microsynth, Switzerland). Editing efficiency was quantified using ICE analysis (Synthego, ICE v3.0).

### Plasmids and engineered cell lines

For in vivo experiments, 1G4 TCR-transduced T cells were generated using the lentiviral vector pCDH-EF1a-1G4 TCR_T2A_copGFP. For in vitro experiments, the corresponding fluorescent reporter construct pCDH-EF1a-1G4 TCR_T2A_mRFP was used. NY-ESO-1-expressing target cells were generated using the plasmid CD811A_PGK_NeoR_EF1a-NYESO full length. This construct was used to generate A375 and Huh7 target cells expressing full-length NY-ESO-1. For Huh7 spheroid experiments, an additional HLA-A*02:01/β2-microglobulin construct provided by A. Lanzavecchia was used to generate cells expressing an EF1a-driven β2-microglobulin–GGGS–HLA-A*02:01 dimer. The Huh7 spheroid target line used in this study lacked endogenous B2M and expressed the HLA-A*02:01–β2-microglobulin dimer, full-length NY-ESO-1, and nuclear GFP.

### Microfluidic device fabrication

Microfluidic devices for water-in-oil (W/O) droplet generation, double-emulsion (DE) conversion, and fluorescence-activated droplet sorting (FADS) were fabricated in-house using standard conventional-and soft-lithography in polydimethylsiloxane (PDMS). Two-dimensional channel layouts were designed in AutoCAD (Autodesk) and transferred to thin-film photolithography masks by an external provider (Micro Lithography Services, Chelmsford, UK). SU-8 3025 photoresist (MicroChem, Westborough, USA) was spin-coated onto silicon wafers, exposed to UV radiation through the mask, and developed to yield molds with channel heights of 33 ± 1 μm for W/O droplet generation devices and 43 ± 1 μm for DE conversion devices.

To facilitate PDMS release, molds were silanized with chlorotrimethylsilane vapor (Sigma-Aldrich) for 2 h. PDMS base and curing agent (Sylgard 184, Dow Corning) were mixed at 10:1 (w/w), degassed, poured over the molds, and cured at 65 °C for 3 h. Cured PDMS slabs were peeled from the molds, cut into individual devices, and punched with 1 mm inlet and outlet ports using a disposable biopsy punch (KAI Medical, Japan). Devices were bonded to glass microscope slides (Thermo Scientific) by oxygen plasma treatment (Zepto Model 1, Diener Electronic). W/O droplet generation devices were rendered hydrophobic by overnight baking at 120 °C. DE conversion devices were rendered permanently hydrophilic by immediate flushing with 1% (w/w) polyvinyl alcohol (molecular weight 13,000–23,000; Sigma-Aldrich) in isopropanol, incubation for 5 min, removal of excess solution, and curing at 120 °C for 15 min. FADS chips were fabricated as previously described by Jain et al. (2024).

### Droplet-based single-cell killing assays and DE conversion

For droplet-based killing assays, JeKo-1 cells were labeled with CellTrace Violet (Thermo Fisher Scientific, cat. no. C34557) according to the manufacturer’s instructions, resuspended at 10^6 cells/ml in cRPMI, and incubated with blinatumomab (2 ng/ml; InvivoGen) for 30 min at 37 °C. Cells were then washed and resuspended at 1.2 × 10^7 cells/ml in cRPMI supplemented with 8 mM CaCl₂ and SYTOX Green Dead Cell Stain (Thermo Fisher Scientific, cat. no. S34860; 1:5,000).

Primary human CD8⁺ T cells were stained with anti-CD2 (1:80; BD Biosciences, cat. no. 555327) or anti-CD8 (1:80; BD Biosciences, cat. no. 561945) in MACS buffer (PBS, 2 mM EDTA, 2.5% FBS) for 10 min at 37 °C, washed, and resuspended at 10^7 cells/ml in cRPMI supplemented with SYTOX Green (1:5,000) and 2 mM EDTA to reduce aggregation.

Cell suspensions were loaded into 500 μl Gastight syringes (Hamilton), and HFE-7500 fluorinated oil containing 2% (w/w) 008-FluoroSurfactant (RAN Biotechnologies, USA) was loaded into a 5 ml Gastight syringe. Syringes were connected to the microfluidic chip using Tygon tubing and 27G blunt needles. Fluids were delivered using neMESYS syringe pumps (CETONI). For co-encapsulation, oil was infused at 10 μl/min and each aqueous cell stream at 3 μl/min. Droplets were collected in 1.5 ml tubes at 30 min intervals and incubated at 37 °C for 1.5, 3, or 6 h. A small vent hole was introduced into each tube cap to permit gas exchange.

For DE conversion, incubated W/O droplets were transferred into a 1 ml syringe and left upright for 5 min to allow droplet accumulation at the syringe tip before injection into the DE conversion chip. The outer aqueous phase consisted of PBS containing 2% (w/w) Poloxamer 188 and 1% (w/w) Tween 20. W/O droplets were injected at 2 μl/min and the outer aqueous phase at 6–8 μl/min to generate DE droplets. DEs were analyzed on a FACSymphony A5 flow cytometer (BD Biosciences), and 100,000 events were acquired per sample.

### Pooled CRISPR screens in microfluidic droplets, FADS, and sample recovery

For pooled screens, JeKo-1 cells were labeled with CellTrace Far Red (Thermo Fisher Scientific, cat. no. C34564), incubated with blinatumomab as above, and resuspended at 1.2 × 10^7 cells/ml in phenol red-free cRPMI supplemented with IL-7 (10 ng/ml), IL-15 (10 ng/ml), SYTOX Green (1:3,500), and CaCl₂ (8 mM). T cells were resuspended at 10^7 cells/ml in phenol red-free cRPMI supplemented with IL-7 (10 ng/ml), IL-15 (10 ng/ml), SYTOX Green (1:3,500), and EDTA (2 mM).

Cells were co-encapsulated as described above. W/O droplets were collected for 40 min in 1.5-ml tubes, incubated at 37 °C for 90 min, and then loaded into a pressure-controlled FADS instrument built in-house at ETH Zurich. The sorting chip contained a reinjection inlet for droplets, two oil inlets to space droplets before sorting, and a Y-shaped junction serving as the sorting point. Sorting was performed at rates between 700-1,000 droplets per second.

Droplets were analyzed on the basis of CellTrace Far Red and SYTOX Green fluorescence. Droplets positive for both CellTrace Far Red and SYTOX Green were sorted as the positive fraction and collected into 1.5-ml tubes containing 300 μl PBS. All remaining droplets were collected separately as the negative fraction.

To recover cells, sorted droplets were destabilized by exposure to a plasma globe, which disrupted the water-in-oil interface and allowed separation of the aqueous phase. The aqueous phase was transferred to low-binding 1.5-ml tubes. For negative or unsorted fractions, excess oil was removed, PBS was added, and samples were processed in the same way. Recovered cells were pelleted at 800 × g for 5 min and lysed in buffer containing 1% SDS, 50 mM Tris, and 10 mM EDTA (pH 8.0). Lysates were stored at -20 °C until genomic DNA extraction.

### Genomic DNA extraction and sgRNA amplification for next-generation sequencing

For genomic DNA extraction, 400 μl lysate aliquots were transferred to 1.5-ml tubes. NaCl (5 M; 16 μl) was added to each tube, and samples were incubated overnight at 66 °C. RNase A (10 mg/ml; 8 μl) was then added and samples were incubated for 1 h at 37 °C, followed by proteinase K (20 mg/ml; 8 μl) for 1 h at 55 °C.

For organic extraction, an equal volume of lysate was mixed with phenol:chloroform:isoamyl alcohol (25:24:1), inverted vigorously, and centrifuged at maximum speed. The aqueous phase was transferred to fresh tubes, and DNA was precipitated with isopropanol, sodium acetate, and GlycoBlue. Samples were frozen at -80 °C until solid, centrifuged at maximum speed for 45 min at 4 °C, washed with 70% ethanol, air-dried, and resuspended in nuclease-free water. DNA concentration was measured using a NanoDrop spectrophotometer.

For PCR1, genomic DNA was split across 50 μl reactions containing at most 3 μg DNA per tube. Each reaction contained Q5 High-Fidelity Master Mix (NEB, cat. no. M0492S), P5 stagger primer mix, and P7 primer. Cycling conditions were 98 °C for 3 min, followed by 25 cycles of 98 °C for 15 s, 57 °C for 15 s, and 72 °C for 25 s, with a final extension at 72 °C for 2 min. PCR2 was used to add Illumina indices and consisted of Q5 High-Fidelity Master Mix, P5 common primer, P7 index primer, and 1:100 diluted PCR1 product in a 25 μl reaction volume. Cycling conditions were 98 °C for 1 min, followed by 12 cycles of 98 °C for 10 s, 66 °C for 10 s, and 72 °C for 30 s, with a final extension at 72 °C for 2 min. All primers were synthesized by Microsynth.

PCR products were gel-purified using a Monarch DNA Gel Extraction Kit (NEB, cat. no. T1020) and further cleaned using SPRI beads (Beckman Coulter). Libraries were sequenced on an Illumina MiSeq instrument using single-end reads.

### CRISPR screen analysis

Sequencing reads were processed using MAGeCK (version 0.5.9.2). sgRNA counts were generated with the MAGeCK count function, normalized using MAGeCK’s median ratio method, and imported into R (version 4.3 or later) for downstream analysis.

For each screen, guides with fewer than 100 normalized counts in either the positive (Pos) or negative (Neg) droplet fraction were excluded. Extreme outliers in the Pos fraction were removed when counts exceeded the screen median by more than 10–15-fold, with the exact threshold adjusted per screen to account for differences in sort purity. For Screen 1, in which sorting of the Pos fraction was inefficient, the Input/Neg ratio was used as a surrogate for killing efficiency. For all subsequent screens, the Pos/Neg ratio was used. Log₂ fold changes (LFCs) were calculated per guide using a pseudocount of 1: log₂((Pos + 1)/(Neg + 1)). LFC values were z-scored within each screen to account for inter-screen variability. Gene-level scores were derived as the median z-scored LFC across all sgRNAs targeting each gene.

Results from 14 independent screens were integrated by joining gene-level median z-LFC values across screens and subtracting each screen’s non-targeting control (NTC) median z-LFC to normalize for baseline shifts. A final median normalized z-LFC was computed per gene across all screens. Statistical significance was assessed by two-sided Welch’s t-test comparing each gene’s per-screen values against the NTC distribution, with the minimum and maximum NTC values removed before testing to reduce the influence of outlier screens. Genes not expressed in T cells (GNG13, CATSPER4, CABYR) were excluded from downstream analysis. JAK1 was excluded as a false positive following manual inspection of individual sgRNAs, which revealed non-specific T-cell depletion rather than a genuine killing phenotype. Statistical analysis and visualization were performed in R using RobustRankAggreg, dplyr, purrr, ggplot2, ggrepel, ggpubr, ggh4x, and plotly.

### Arrayed validation assays

Primary human memory CD8⁺ T cells were edited by RNP electroporation as described above and expanded for 10 days before functional testing.

For droplet-based validation, gene-edited and NTC T cells were co-encapsulated with JeKo-1 cells under the same conditions described for the droplet assay. To minimize batch effects, edited and control cells were processed in parallel within the same encapsulation run. Droplets were collected twice at 30 min intervals. One batch was immediately converted into DEs and analyzed as the 0 h time point, whereas the second batch was incubated at 37 °C for 90 min before DE conversion. Both samples were analyzed by flow cytometry to determine target-cell killing.

For bulk killing assays, JeKo-1 cells were labeled with CFSE (BioLegend, cat. no. 423801), resuspended at 10^6 cells/ml in cRPMI, and incubated with blinatumomab (2 ng/ml) for 30 min at 37 °C. Cells were then plated in 96-well plates in 50 μl cRPMI per well. Edited T cells were added at effector-to-target ratios of 1:3, 1:1, and 3:1, and cRPMI was added to a final volume of 200 μl. Each condition was prepared in triplicate or quadruplicate as indicated. At 0, 4, 8, and 24 h, plates were centrifuged at 450 × g for 5 min. Supernatants were collected and stored at -20 °C, and cell pellets were resuspended in MACS buffer containing DAPI (1:200; BD Biosciences, cat. no. 564907) for flow-cytometric analysis of cell death.

For solid-tumor killing assays, A375 cells were transduced to express NY-ESO-1 (A375^OE). Primary CD8⁺ and CD4⁺ T cells were transduced with a lentiviral construct encoding the 1G4 NY-ESO-1 TCR linked to RFP by a T2A sequence. One day later, T cells were edited by RNP electroporation. Three days after electroporation, 1G4 TCR expression was assessed by RFP positivity. A375^OE cells were seeded at 20,000 cells per well in flat-bottom 96-well plates. Edited T cells were added at defined effector-to-target ratios based on the percentage of RFP⁺ cells. Plates were imaged every 4 h for 3 days using an IncuCyte live-cell imaging system. At the final time point, cells were harvested and resuspended in MACS buffer containing DAPI for flow-cytometric analysis of cell death.

For Huh7 spheroid assays, Huh7 target cells lacking endogenous B2M and expressing the HLA-A*02:01–β2-microglobulin dimer, full-length NY-ESO-1, and nuclear GFP were seeded at 5,000 cells per well in Nunclon Sphera 96-well U-bottom low-attachment plates (Thermo Fisher Scientific, cat. no. 174925) in cDMEM and incubated for 48 h to allow spheroid formation. 1G4 TCR-transduced T cells were then added at an effector-to-target ratio of 3:1, calculated on the basis of TCR-positive (RFP-positive) cells, in 100 μl cRPMI without exogenous cytokines. Plates were transferred to an IncuCyte live-cell imaging system (Sartorius), and images were acquired every 4 h using brightfield, green, and red channels with a 4× objective for up to 6 days. Images were analyzed using IncuCyte software version 2024A.

### Rapamycin treatment

For in vitro mTORC1 inhibition experiments against JeKo-1 cells, bulk CD8⁺ T cells were isolated from healthy donors and activated for 2 days on plate-bound anti-CD3 and anti-CD28 antibodies. Cells were then expanded for 10 days in cRPMI supplemented with IL-7 (10 ng/ml) and IL-15 (10 ng/ml). Where indicated, cells were treated with rapamycin (5 nM) for 8 days before challenge with JeKo-1 cells. Cytotoxicity was then assessed as described for bulk killing assays.

For in vivo manufacturing experiments, rapamycin was added at 20 nM for 8 days during the expansion phase as described below.

### Mass spectrometry sample preparation

Primary human CD8⁺ T cells carrying RPTOR or RHEB knockout were generated as described above. After 10 days of expansion, 3 × 10^6 resting T cells were collected and snap-frozen. For each condition, technical triplicates were generated by dividing samples into 1 × 10^6 cells per replicate. For activated conditions, 3 × 10^6 cells were stimulated for 3 days with anti-CD3 and anti-CD28 before snap-freezing.

Cell pellets were lysed in 8 M urea prepared in 50 mM ammonium bicarbonate and sonicated for 15 min in a Bioruptor (15 cycles of 30 s on/30 s off). Proteins were reduced with 10 mM dithiothreitol for 20 min at room temperature and alkylated with 50 mM iodoacetamide for 30 min in the dark. Samples were first digested with LysC (1:100 enzyme-to-protein ratio) for 2 h at room temperature, diluted to 2 M urea with 50 mM ammonium bicarbonate, and then digested overnight with sequencing-grade modified trypsin (1:100 enzyme-to-protein ratio). Digestion was stopped by addition of 2% acetonitrile and 0.3% trifluoroacetic acid. Debris was removed by centrifugation, and peptides were desalted on C18 StageTips. Eluted peptides were dried in a SpeedVac and resuspended in 2% acetonitrile, 0.5% acetic acid, and 0.1% trifluoroacetic acid prior to mass spectrometry analysis.

### Proteomics data analysis

Raw DIA-MS/MS data were quantified using DIA-NN and exported as protein-group intensity matrices. Two independent proteomics experiments were analyzed. For the differential abundance analysis shown here, resting NTC and RPTOR-KO samples from both experiments were combined.

Protein intensities corresponding to multiple isoforms with the same gene symbol were summed before analysis. Summed intensities were log₂-transformed, and zero values were treated as missing. To correct for between-run batch effects introduced by the two acquisitions, log₂ intensities were adjusted using removeBatchEffect from the limma package, with experimental group included in the design matrix.

Differential abundance testing was performed using repeated-imputation t-tests under a missing-not-at-random assumption. Missing values were imputed 10 times independently by sampling from a downshifted normal distribution (downshift 1.8 SD; width 0.3 SD of the observed values per sample). A two-sided Welch’s t-test was performed for each imputed dataset, and P values and log₂ fold changes were averaged across imputations. Multiple-testing correction was applied using the Benjamini-Hochberg method. Proteins with an absolute log₂ fold change greater than 1 and a false discovery rate below 0.05 were considered significantly changed.

Estimated protein copy numbers per cell were calculated from raw intensities using total-protein-mass normalization, assuming a resting T-cell protein content of 35 pg per cell. Molecular weights were obtained from reviewed human UniProt Swiss-Prot entries. Copy number was calculated as:

copy number = (protein intensity / total intensity) × cell mass / molecular weight × Avogadro’s constant

All analyses were performed in R (version 4.3 or later) using limma, ggplot2, ggrepel, ggpubr, and ggh4x.

### LC-MS/MS analysis of metabolites

The quantitative measurement of amino acids was performed on a QTRAP® 6500+ (Sciex, MA, USA) coupled to a Waters Acquity® UPLC system (MA, USA). The chromatographic separation was performed using an Acquity® Premier BEH Amide column (2.1 ×100 mm, 1.7 µm; Waters) at a temperature of 30°C in conjunction with a gradient of mobile phases A (20 mM NH4HCO3 in water; pH 9) and B (20 mM NH4HCO3 in 90% Acetonitrile; pH 9). A CE-IVD-validated commercial product (Chromsystems, Germany; Product 75111) for the quantitative measurement of amino acids was used and optimised for cell lysates. Ten microliters of reconstituted calibrators, quality control (QC) samples and sample material, all kept at 4°C in the autosampler, were injected into the ultra-performance liquid chromatography-tandem mass spectrometry (UPLC-MS/MS) system at a flow rate of 0.5 milliliters per minute, including 90% mobile phase B. Following a 1-minute period, the mobile phase A started to linearly increase up to 35% for 4 minutes, after which it was held for 1 minute. The gradient returned to the initial conditions within 0.5 minutes, followed by a further 3.5 minutes for column equilibration. Analytes were detected in multiple reaction monitoring mode (MRM) using positive electrospray ionization (ESI).

### Immunoblotting

T cells transduced with either a non-targeting control (NTC) gRNA or an RPTOR-targeting gRNA were seeded at (4 \times 10^6) cells/mL in 50 μL of RPMI medium in wells coated with anti-CD3 and anti-CD28 antibodies. Cells were stimulated at 37°C for 0, 5, 15, or 30 min. Following stimulation, cells were washed twice with ice-cold PBS and lysed directly in 25 μL of Laemmli sample buffer per well containing 187.5 mM Tris-HCl (pH 6.8), 6% SDS, 0.03% bromophenol blue, and 30% glycerol.

Lysates were heated at 95°C for 5 min, and proteins were separated by SDS–PAGE on precast 4–20% gradient gels (GenScript) using MOPS running buffer. Proteins were transferred to nitrocellulose membranes, which were blocked for 30 min at room temperature in 5% non-fat dry milk prepared in PBS containing 0.1% Tween-20 (PBST). Membranes were incubated overnight at 4°C with primary antibodies against phospho-AKT (Ser473; clone 4E2, Cell Signaling Technology, #9271S), phospho-p70 S6 kinase (Thr389; Cell Signaling Technology, #9205), total p70 S6 kinase (Cell Signaling Technology, #9202), and β-actin (clone E4D9Z, Cell Signaling Technology, #58169), diluted in 5% non-fat dry milk in PBST.

Membranes were washed three times with PBST and incubated for 30 min at room temperature with the appropriate HRP-conjugated secondary antibodies (Cell Signaling Technology; 1:2,500). Following three 10-min washes with PBST, chemiluminescent signals were developed using WesternBright ECL HRP or WesternBright Quantum substrates (Advansta). Images were acquired using a Fusion FX imaging system (Vilber), and band intensities were quantified using ImageJ.

### Preparation of 1G4 TCR-transduced T cells for in vivo experiments

CD4⁺ and CD8⁺ T cells were isolated from healthy donor buffy coats, mixed at a 1:1 ratio, and activated on plate-bound anti-CD3 and anti-CD28 antibodies in cRPMI supplemented with IL-7 (10 ng/ml) and IL-15 (10 ng/ml). Twenty-four hours later, cells were transduced with 1G4 TCR lentivirus by spin infection at 800 × g for 1 h at 32 °C. Cells were removed from plate-bound stimulation the following day, corresponding to a total of 48 h of activation, and expanded for 10 days in cRPMI supplemented with IL-7 (10 ng/ml), IL-15 (10 ng/ml), and IL-2 (50 U/ml). Where indicated, cells were treated with rapamycin (20 nM) for 8 days during the expansion phase.

### In vivo tumor model

All animal experiments were performed under authorization number 38308. NSG mice aged 8–12 weeks were injected subcutaneously in the right flank with 1 × 10^6 A375 cells expressing full-length NY-ESO-1 in 50 μl PBS. After 7–8 days, when tumors reached approximately 20–40 mm^3^, mice received intravenous injection of 1G4 TCR-transduced T cells in 100 μl PBS. T-cell doses were normalized on the basis of the percentage of TCR-positive cells to approximate an effector-to-target ratio of 1:1 relative to tumor cells, considering only TCR-positive cells. In the experiment shown, the total number of infused cells was 4.2 × 10^6^ per mouse for the untreated 1G4 T-cell group and 4.8 × 10^6^ per mouse for the rapamycin-treated 1G4 T-cell group, corresponding in each case to 1 × 10^6^ TCR-positive cells. Tumor growth was monitored three times per week using caliper measurements.

