## Supplemental Material for "Droplet single-cell CRISPR screens identify regulators of T cell–mediated target-cell killing"

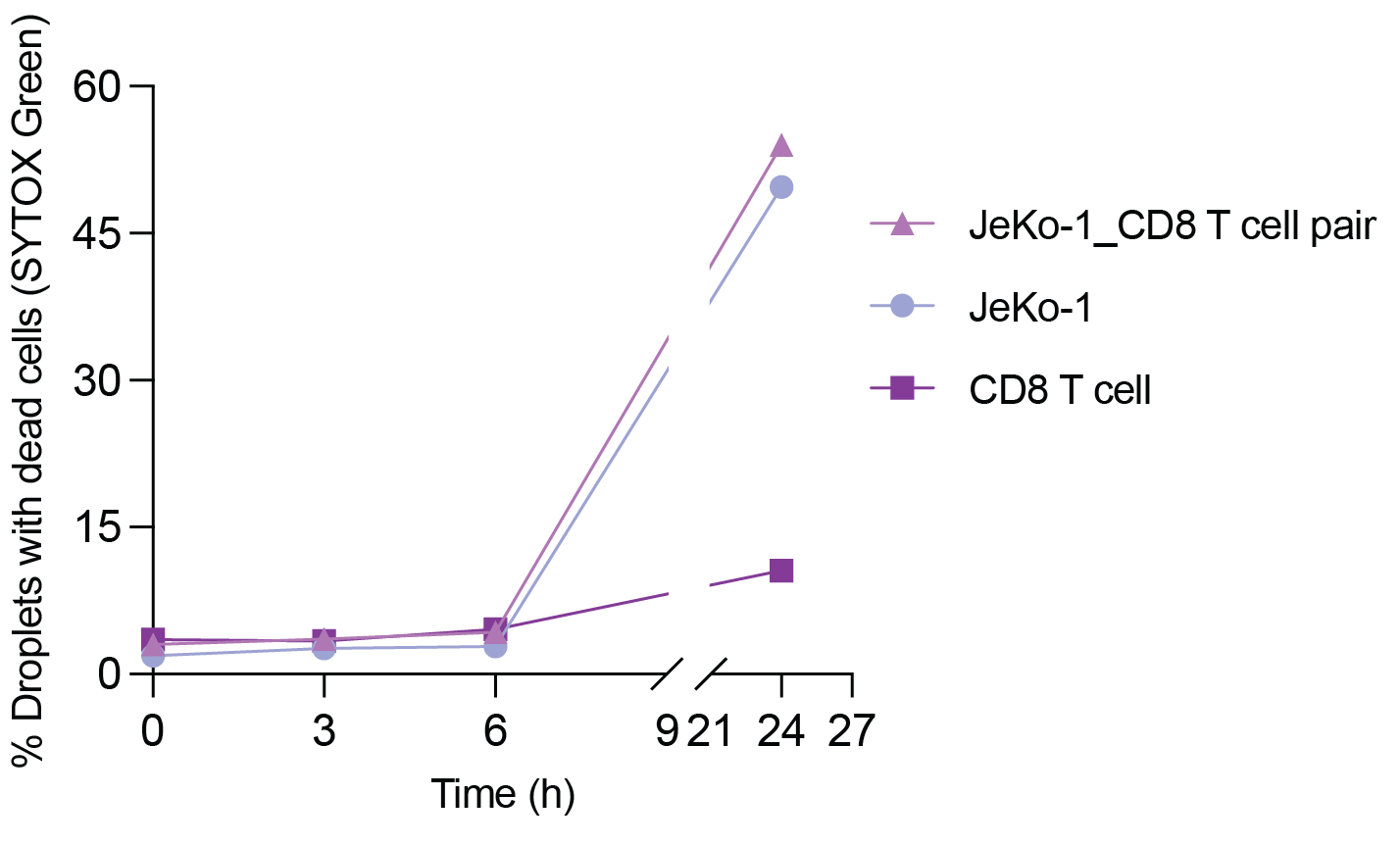

**Suppl. Fig. 1. T cells and JeKo-1 cells remain viable in droplets for greater than 6 hours**

Primary human CD8⁺ T cells, CD19⁺ JeKo-1 lymphoma cells, or paired CD8⁺ T cell–JeKo-1 droplets were generated in the absence of blinatumomab, converted to double emulsions, and analyzed by flow cytometry at the indicated time points. The percentage of droplets containing SYTOX Green-positive cells remained low for all conditions over the 6-h assay window, indicating that T cells and JeKo-1 cells are largely viable in droplets under control conditions. At later time points (>24 h), JeKo-1 viability declined in double-emulsion droplets.

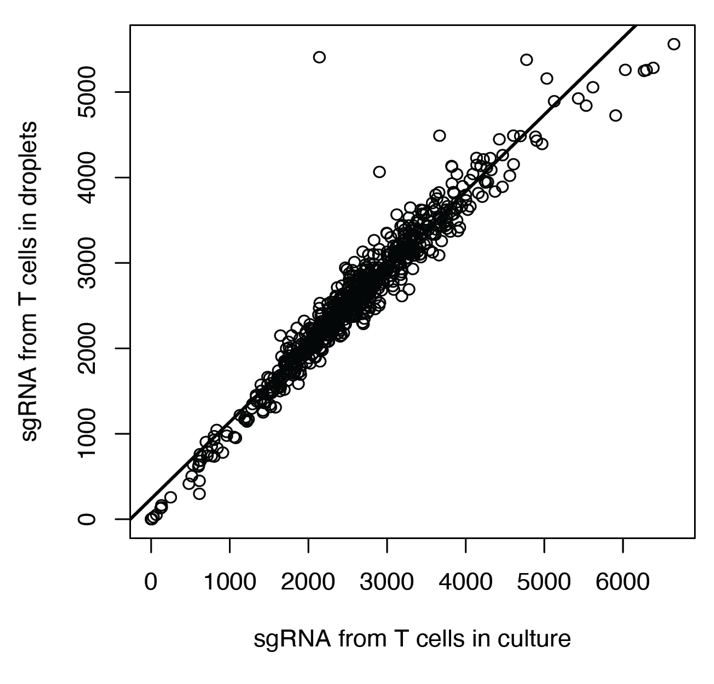

**Suppl. Fig 2. Efficient recovery of sgRNAs from T cells encapsulated into droplets**

sgRNA abundance measured by next-generation sequencing in T cells maintained in culture was compared with sgRNA abundance recovered from T cells encapsulated in droplets and processed through the droplet workflow. Each point represents one sgRNA, and the line indicates the linear regression fit. The strong concordance between conditions indicates efficient sgRNA recovery and preservation of library representation following droplet encapsulation and processing (R² = 0.93).

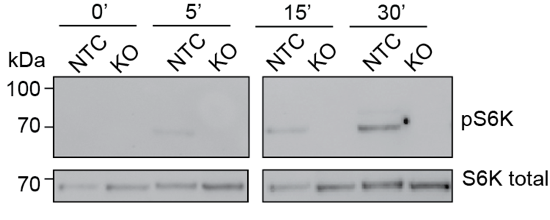

**Suppl. Fig. 3 Knockout of *RAPTOR* diminishes phosphorylation of S6K**

Representative immunoblot of phospho-pS6K (Thr389) in NTC and RPTOR-KO CD8⁺ T cells following anti-CD3/CD28 stimulation for the indicated times. Total S6K served as a loading control.

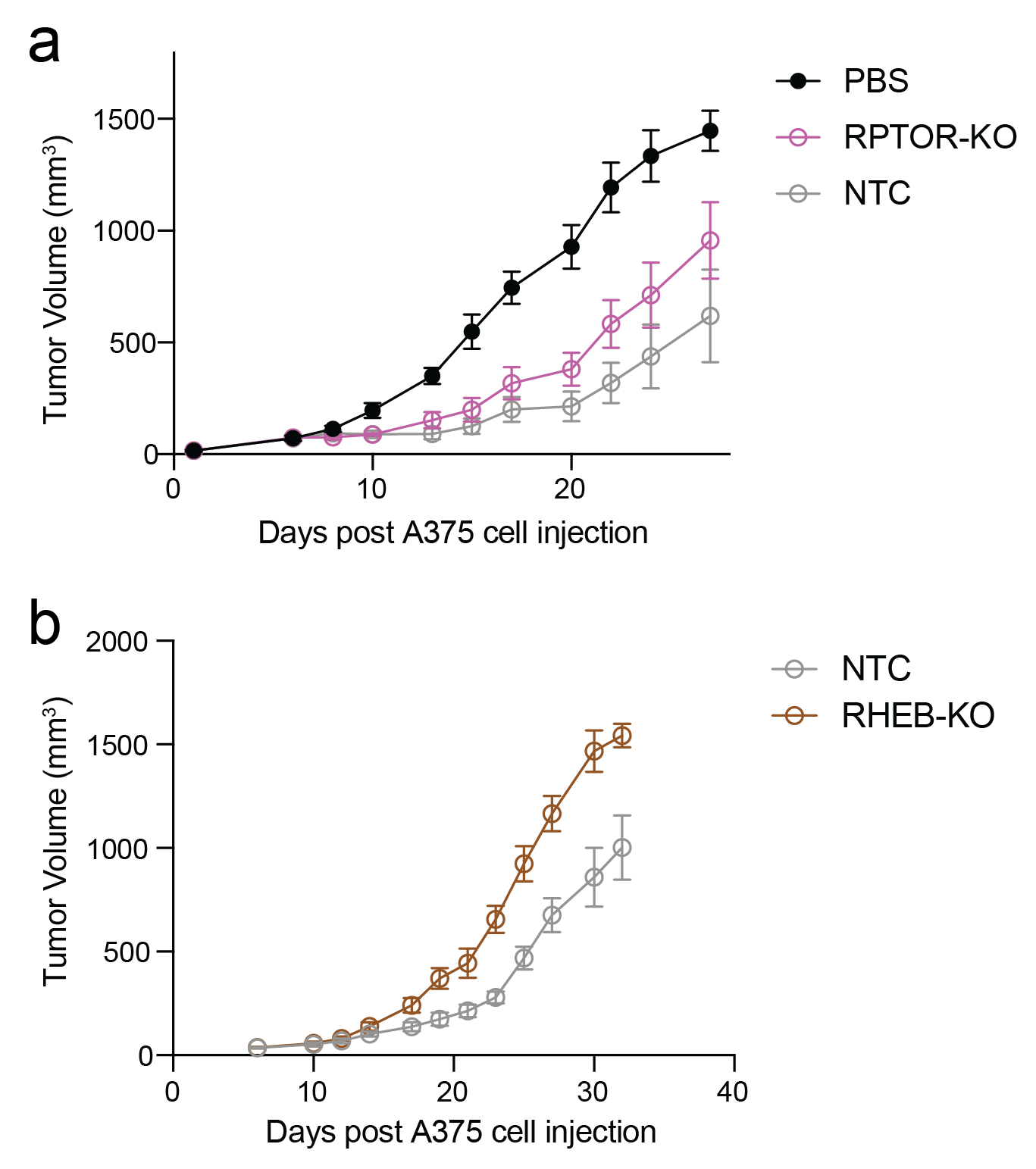

**Suppl. Fig. 4. Knockout of RPTOR and RHEB diminishes antitumor activity in vivo**

**(a**) Tumor growth in A375 tumor-bearing NSG mice treated with PBS, 1G4 TCR-transduced T cells treated with a non-targeting control sgRNA (NTC), or 1G4 TCR-transduced T cells in which *RPTOR* was ablated. Data are mean tumor volume ± SEM; n = 10 (PBS), 10 (NTC), and 10 (*RPTOR*-KO).

**(b**) Tumor growth in A375 tumor-bearing NSG mice treated with 1G4 TCR-transduced T cells treated with a non-targeting control sgRNA (NTC), or 1G4 TCR-transduced T cells in which *RHEB* was ablated. Data are mean tumor volume ± SEM; n = 10 (NTC), and 8 (*RHEB*-KO).

**Tables:**

Suppl. Table 1. sgRNA library used for the CRISPR screens

Suppl. Table 2. Gene-level log fold changes for 14 CRISPR screens. Each screen was performed with primary human CD8⁺ T cells from a different donor.
